# Parallel Contributions of Externalizing Polygenic Liability and Brain Imaging Phenotypes to Adolescent Substance Use Initiation Timing: A Multistage Analysis in the ABCD Study

**DOI:** 10.64898/2026.04.08.717299

**Authors:** Mengman Wei, Qian Peng

## Abstract

**Background:** Adolescent substance use initiation is shaped by multiple genetic and neurobiological factors. Externalizing liability, a transdiagnostic genetic dimension capturing shared predisposition to impulsivity, disinhibition, and related traits, is among the strongest polygenic predictors of early substance initiation. Yet how this genetic risk relates to brain structure and function, and whether baseline brain phenotypes statistically account for or instead act in parallel with genetic liability, remains unresolved.

**Methods:** Using the ABCD Study, we analyzed an analytic cohort of 10,608 participants with genotype data, baseline multimodal neuroimaging-derived phenotypes (IDPs), and longitudinal substance initiation assessments. Outcome-specific models included up to 10,599 participants after complete-case filtering for survival variables and covariates. We implemented a multistage framework linking an externalizing polygenic risk score (extPRS) to baseline IDPs and longitudinal substance initiation outcomes, including alcohol, nicotine, cannabis, and any substance. Stage 1 screened extPRS–IDP associations using covariate-adjusted linear models with false discovery rate (FDR) control. Stage 2 estimated extPRS effects on time-to-initiation using Cox proportional hazards models. Stage 3 fit joint extPRS + IDP Cox models to identify IDPs that predicted initiation beyond extPRS. Stage 4 conducted bootstrap-based mediation analyses to quantify average causal mediation effects (ACME), average direct effects (ADE), and the proportion of the extPRS–initiation association statistically accounted for by individual IDPs.

**Results:** Higher extPRS was robustly associated with earlier initiation across all substances: alcohol, hazard ratio (HR) = 1.13; nicotine, HR = 1.63; cannabis, HR = 1.67; and any substance, HR = 1.15. Thousands of extPRS-associated IDPs were identified at baseline, with highly concordant effect profiles across robustness specifications. In joint models, numerous IDPs independently predicted initiation timing above and beyond extPRS: 31 for alcohol, 32 for any substance, 137 for cannabis, and 459 for nicotine, with a replicated core set across specifications. Cannabis and nicotine initiation were jointly predicted by superficial white matter (SWM) microstructural integrity in sensorimotor cortex as a protective factor, and by irregular activity in a right-hemisphere region as a risk factor. Alcohol initiation was predicted by a largely distinct, strongly left-lateralized frontolimbic SWM intensity axis. Nicotine initiation additionally and uniquely involved restricted gray matter diffusion in the anterior cingulate cortex and subcallosal cortex. Despite these robust independent IDP associations, mediation analyses showed that indirect effects through individual baseline IDPs were very small in magnitude (ACME ≈ 10^−4^), accounting for less than 2% of the total extPRS effect, with FDR-significant mediation surviving only for alcohol and any-substance initiation.

**Conclusions:** Within the scope of externalizing polygenic risk and baseline neuroimaging, the predominant pattern is one of largely parallel, additive contributions to adolescent substance initiation rather than a dominant genetic → brain → behavior pathway. Baseline brain features, particularly prefrontal functional variability and frontolimbic and sensorimotor white matter integrity, predict initiation risk beyond extPRS, indicating neurobiological vulnerabilities not captured by this genetic dimension. However, these baseline IDPs explain only a small fraction of the extPRS–initiation association, suggesting that externalizing genetic liability may operate through pathways not fully represented by cross-sectional baseline imaging. Whether other genetic risk dimensions, such as substance-specific PRS, or dynamic longitudinal brain measures show stronger mediation patterns remains an important open question.

## Introduction

Substance use initiation during adolescence is a major public health concern. The majority of individuals who will ever develop a substance use disorder (SUD) begin using before the age of 18, and earlier age of onset is among the strongest predictors of problematic use in adulthood [1, 2, 3, 4, 5, 6, 7]. National surveillance data indicate that by the end of high school, a substantial minority of adolescents have tried alcohol, cannabis, or tobacco, with initiation rates rising sharply across early-to-mid adolescence [8]. Identifying the biological and psychosocial factors that confer vulnerability to early initiation is therefore a priority for prevention and intervention.

Substance use initiation is moderately to substantially heritable, with twin and family studies estimating heritability in the range of 40–70% depending on the substance and population studied [9, 10]. Large-scale genome-wide association studies (GWAS) have now characterized the genetic architecture of individual substances, including alcohol use disorder and consumption traits [11], cigarette smoking [12], cannabis use [13], and problematic substance use more broadly [14]. Crucially, genetic associations across substances are not independent: liability to different forms of externalizing psychopathology, including antisocial behavior, substance use disorders, ADHD, and related conditions, shares a substantial genetic basis captured by an externalizing spectrum [1, 15]. The externalizing GWAS underlying the polygenic risk score used in the present study was derived from a multivariate analysis of over 1.5 million individuals, capturing this transdiagnostic genetic dimension with high reliability and generalizability [15]. This externalizing PRS (extPRS) captures genetic liability to the broad spectrum of disinhibited behavior rather than the specific genetic architecture of any single substance, and it is distinct from substance-specific PRS, such as those for alcohol use disorder, nicotine dependence, or cannabis use disorder, that may capture more specific genetic influences on individual substances. The relationships between these different genetic dimensions, and the degree to which each relates to brain structure and function, are only beginning to be characterized.

A prominent neurodevelopmental hypothesis holds that adolescent risk behavior, including substance initiation, arises from a transient developmental imbalance: reward and motivational systems mature earlier than the prefrontal control systems that regulate them, creating a window of heightened impulsivity and susceptibility to environmental incentives [16, 17, 18]. This dual-systems model has motivated a large body of imaging research examining whether structural or functional differences in frontal–subcortical circuitry precede and predict the onset of substance use [19, 20]. Prior studies have documented associations between cortical thickness, white matter microstructure, and resting-state connectivity and concurrent or subsequent substance use in adolescent samples [21, 22], and the launch of large-scale prospective cohorts, especially the Adolescent Brain Cognitive Development (ABCD) Study, has substantially expanded the statistical power available to test these relationships [23, 24, 25, 26, 27]. Baseline imaging data from the ABCD Study have revealed that structural and functional brain phenotypes measured at ages 9–10 are associated with substance initiation and related risk behaviors, even after controlling for sociodemographic factors [28]. However, the mechanisms through which these imaging phenotypes relate to genetic risk and to substance initiation timing remain incompletely understood.

A natural question is whether genetic liability to externalizing behavior influences substance initiation through its effects on brain development, consistent with a mediation model, or whether genetic risk and brain phenotypes operate as largely parallel contributors to initiation risk through partially distinct mechanisms. Under the mediation model, the extPRS would be expected to influence initiation primarily by shaping neurodevelopmental outcomes, such as altered frontal maturation, white matter organization, or network dynamics, that in turn lower the threshold for substance use. Under a parallel-pathways model, extPRS would influence initiation via biological routes not well captured by baseline brain structure and function, such as neurochemical differences, gene expression, peripheral or developmental processes that precede the imaging window, while brain features simultaneously reflect additional and largely independent sources of variance, including environmental exposures, epigenetic factors, fetal development, or genetic influences outside the externalizing spectrum, that also shape initiation risk. Distinguishing these models is nontrivial. It requires large prospective samples, rigorous multiple-testing control, and analytical designs that explicitly test both the independent predictive value of brain phenotypes beyond genetic liability and the proportion of genetic effects that is statistically explained by brain measures. It is also important to recognize that this question does not have a single answer: different genetic dimensions, such as substance-specific PRS versus transdiagnostic externalizing PRS, and different brain measurement models, such as baseline cross-sectional phenotypes versus longitudinal change or resting-state versus task-based activation, may yield quite different conclusions about the degree of mediation or independence.

The present study addresses this question specifically within the scope of externalizing polygenic liability and baseline multimodal neuroimaging, using the ABCD Study’s uniquely large and deeply phenotyped sample. We implemented a four-stage multistage analysis framework that: (i) screens extPRS associations with thousands of imaging-derived phenotypes (IDPs); (ii) characterizes direct extPRS effects on time-to-substance-initiation across four outcomes; (iii) tests whether extPRS-associated IDPs predict initiation beyond extPRS in joint models; and (iv) formally quantifies the proportion of extPRS effects mediated through individual IDPs. This design allows us to distinguish IDPs that merely correlate with both genetic liability and initiation from those that carry independent predictive information, and to precisely estimate the degree to which baseline brain phenotypes mediate versus parallel the genetic signal. Our aims are to: (1) characterize the extent and structure of extPRS–IDP associations across brain modalities; (2) quantify direct extPRS effects on substance initiation timing using survival models; (3) identify brain phenotypes that predict initiation timing independently of extPRS; and (4) estimate the proportion of extPRS effects on initiation attributable to baseline brain IDPs. We interpret findings in light of the specific scope of the extPRS measure and the baseline imaging window, while highlighting what the results imply about the broader landscape of genetic and neural pathways to adolescent substance initiation.

## Methods

### Cohort and outcomes

We included ABCD Study^®^ participants with available genotype data sufficient to compute an externalizing polygenic risk score (EXTPRS), at least one eligible neuroimaging phenotype, and longitudinal substance-initiation assessments. The primary outcome was alcohol initiation; secondary outcomes were initiation of nicotine, cannabis, and any substance.

Time-to-initiation was defined as the interval from baseline assessment to the first study visit at which the participant reported initiation of the relevant substance. Participants without initiation were censored at their last available follow-up visit. We constructed (time, event) variables for Cox proportional hazards models accordingly. EXT-PRS computation and substance-initiation phenotype harmonization followed previously described procedures [29, 30, 28, 15, 31, 32, 33], with key definitions summarized here and full implementation details provided in those reports.

### Externalizing polygenic risk score (EXT-PRS)

We derived an externalizing polygenic risk score (EXT-PRS) using externalizing GWAS summary statistics and ABCD genotype data after standard quality control. Summary statistics were harmonized to the ABCD reference alleles and genomic build as needed, and the PRS was computed using our previously published pipeline [29, 30, 28, 15, 31, 32, 33]. The EXT-PRS was *z*-standardized (mean = 0, SD = 1) prior to regression analyses.

### Neuroimaging phenotypes (IDPs)

We analyzed baseline multimodal neuroimaging-derived phenotypes (IDPs) from the ABCD core imaging tables, including structural MRI (e.g., cortical thickness, surface area, regional volumes, and intensity/contrast metrics), diffusion MRI measures (DTI and restriction spectrum imaging [RSI] summaries), and resting-state fMRI summary measures (e.g., connectivity-and variance-related metrics).

IDPs were extracted at the region-of-interest level across multiple parcellation and segmentation schemes (e.g., aseg and cortical atlas variants). For each table, we retained one baseline record per participant and standardized each IDP (*z*-score) within the analytic sample to improve comparability across modalities and scales.

### Covariates

We examined associations between EXT-PRS and IDPs using standardized linear regression. For each IDP, we fit a linear model with the IDP as the dependent variable and EXT-PRS as the main predictor, adjusting for key demographic and imaging covariates: age, sex, ancestry principal components (PCs), imaging site, scanner manufacturer, and scanner device serial number.

Covariate sets were modality-specific where appropriate. For structural MRI phenotypes, we additionally adjusted for intracranial volume (ICV) and, when applicable, global mean cortical thickness or total surface area. For diffusion MRI and fMRI measures, we further included motion and data-quality indices to account for acquisition-related artifacts.

Covariates were harmonized across modalities, with categorical variables one-hot encoded and all continuous variables winsorized at the 0.5% tails. Models were estimated using ordinary least squares (OLS) in `statsmodels`. Robust standard errors were computed, with clustering by site when specified. Associations were summarized using standardized *β* coefficients, standard errors, *p*-values, and partial *R*^2^ values, where partial *R*^2^ represents the variance uniquely explained by EXT-PRS.

To control for multiple testing across IDPs, we applied both Benjamini–Hochberg false discovery rate (FDR) [34] and Bonferroni correction [35]. In addition, we estimated the effective number of independent tests (*M*_eff_) using the Li and Ji method [36] to account for correlation among IDPs. FDR-significant results after Li– Ji correction were reported separately for each model specification, including site-clustered and SES-adjusted sensitivity analyses.

### Externalizing polygenic risk score (extPRS), neuroimaging phenotypes (IDPs), and substance use initiation: multistage analysis pipeline

We implemented a four-stage analysis pipeline to connect externalizing polygenic liability (extPRS) with baseline neuroimaging-derived phenotypes (IDPs) and time-to-initiation of substance use (alcohol, nicotine, cannabis, and any substance). In brief, we (i) screened extPRS–IDP associations across all IDPs, (ii) tested the direct association of extPRS with initiation using Cox models, (iii) fit joint extPRS + IDP Cox models to prioritize IDPs associated with initiation beyond extPRS, and (iv) evaluated mediation for prioritized extPRS–IDP–initiation candidate pairs.

#### Stage 1: extPRS → IDP association screening (IDP-by-IDP linear regression)

##### Data inputs and preprocessing

The extPRS was computed and standardized to a *z*-score within the analytic sample. Baseline IDPs were loaded from a single harmonized IDP table, and covariates were loaded from a precomputed covariate table.

By default, we analyzed all numeric IDP columns, excluding non-phenotype fields (e.g., unnamed columns). For each IDP, we merged IDP values with extPRS and covariates, removed observations with missing values for model variables, and applied two-sided winsorization to the IDP outcome (default: 0.5% in each tail) to reduce the influence of extreme values. Each IDP was then standardized to a *z*-score within the analytic subset used for that model. IDPs with zero or near-zero variance after preprocessing were excluded.

##### Covariates

All Stage 1 models included the following covariates when available: age, sex, genetic ancestry principal components (PC1–PC20), site (modeled using one-hot encoding with *k* − 1 dummy variables), scanner manufacturer (one-hot encoded), and scanner device serial number (one-hot encoded).

To better control modality-specific confounding, we applied modality-aware covariate rules based on the IDP name:

- For non-T1 modalities (e.g., diffusion MRI or fMRI; identified by keywords such as `dti, dmri, fmri, rsi`), we additionally included a motion covariate when available.
- For structural surface area or volume measures (identified by keywords such as `area, vol, volume`), we additionally adjusted for intracranial volume (ICV) to account for head size.

##### Statistical model

For each IDP, we fit an ordinary least squares regression:

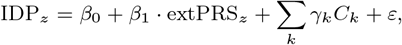

where *C*_*k*_ denotes the covariates defined above. For each IDP, we recorded the sample size, *β*_1_, robust site-clustered and HC3 standard errors, *p*-values, model *R*^2^, and partial *R*^2^ for extPRS.

##### Multiple-testing control

Across all tested IDPs, we controlled for multiple comparisons using (i) Benjamini–Hochberg false discovery rate (BH-FDR) across IDPs and (ii) Bonferroni correction across IDPs.

In addition, to account for correlation among IDPs when defining the discovery “hit” set used in later stages, we applied an effective number of independent tests approach [36]. The resulting discovery-significant IDPs were carried forward to Stage 3.

#### Stage 2: extPRS → initiation (direct effect; Cox proportional hazards models)

##### Outcomes and survival setup

For each initiation outcome (alcohol, nicotine, cannabis, and any substance), we modeled time-to-initiation using a Cox proportional hazards framework, where *time* denotes the follow-up time to initiation or censoring, and *event* indicates initiation (1 = initiated, 0 = censored).

##### Model specification

We fit:

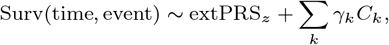

using Efron’s method for ties. Covariates matched the baseline set used in Stage 1 (age, sex, ancestry principal components, and available site/scanner indicators). We used site-clustered standard errors. Results are reported as hazard ratios (HRs) with 95% confidence intervals for extPRS.

#### Stage 3: extPRS + IDP → initiation (joint Cox models; candidate IDP prioritization)

##### Candidate IDPs

We restricted Stage 3 to the discovery-significant IDPs identified in Stage 1 after multiple-testing control, including correlation-aware adjustment.

##### Joint Cox models

For each outcome and each candidate IDP, we fit:

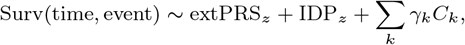

again using Efron’s method for ties and site-clustered standard errors when available. Both extPRS and the IDP were re-standardized within the analytic subset for each model (after missing-data filtering), such that effect sizes correspond to a one standard deviation increase.

##### Multiple-testing control in Stage 3

Within each outcome, we applied Benjamini–Hochberg false discovery rate (BH-FDR) correction across candidate IDPs separately for each term (i.e., one correction for the extPRS term and one correction for the IDP term). Outcome–IDP pairs were retained as mediation candidates when the IDP_*z*_ term satisfied FDR *<* 0.05.

#### Stage 4: Mediation analysis (ACME/ADE; bootstrap inference)

##### Candidate pairs and model forms

For each outcome–IDP pair passing Stage 3, we evaluated whether the IDP statistically mediated the association between extPRS and initiation. We treated extPRS_*z*_ as the exposure and IDP_*z*_ as the mediator, using:

- **Mediator model (linear):**

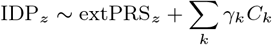

- **Outcome model (logistic):**

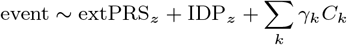

where *event* is the binary initiation indicator (consistent with the implementation used for mediation estimation). Covariates matched those used in earlier stages (age, sex, ancestry principal components, and available site/scanner indicators). extPRS and IDP were standardized within the mediation sample.

##### Quality control thresholds

To reduce unstable estimates, we required:

- at least 200 participants per outcome–IDP mediation analysis,
- at least 20 events and 20 non-events, and
- non-negligible variance in extPRS_*z*_ and IDP_*z*_.

##### Estimation and multiple-testing correction

We estimated the average causal mediation effect (ACME), average direct effect (ADE), and proportion mediated using nonparametric bootstrap inference with a target of 5,000 simulations. If bootstrap inference failed to converge, we used a pre-specified fallback sequence (bootstrap with 1,000 simulations; if needed, quasi-Bayesian simulation with 1,000 draws).

Finally, within each outcome we applied BH-FDR correction to mediation *p*-values (for ACME and proportion mediated).

## Results

### Sample characteristics

The analytic cohort included 10,608 ABCD participants with genotype data sufficient to compute the externalizing polygenic risk score (EXT-PRS), baseline neuroimaging phenotypes, and required covariates, and thus contributed to all initiation analyses.

Baseline age was 9.48 years (SD = 0.51; median = 9, IQR = 1). The sex distribution was 5,078 male (47.9%), 4,650 female (43.8%), and 880 unknown (8.3%). Ancestry composition was 5,710 White (53.8%), 2,077 Hispanic (19.6%), 1,480 Black (14.0%), 1,116 Other (10.5%), and 225 Asian (2.1%).

During follow-up, initiation events were observed for alcohol (*n* = 3,964; 37.4%), nicotine (*n* = 591; 5.6%), cannabis (*n* = 378; 3.6%), and any substance (*n* = 4,302; 40.6%), with remaining participants censored at their last available visit.

### Stage 1: extPRS → IDP association screening (IDP-by-IDP linear regression)

Using our primary specification (site-clustered standard errors with SES covariates included), we identified 3,307 IDPs significantly associated with extPRS (Li–Ji-adjusted BH-FDR *q* ≤ 0.05). Sensitivity analyses showed robust findings across alternative specifications, including heteroskedasticity-robust (HC3) standard errors and exclusion of SES covariates. Specifically, we identified 5,460 hits for HC3 with SES included, 5,419 for site-clustered SE without SES, and 4,641 for HC3 without SES.

Despite differences in total discoveries, signals were highly stable across specifications: 3,184 IDPs were significant in both SES-on models, and 3,098 IDPs remained significant across all four settings (SES on/off × site-clustered/HC3). Effect sizes were highly concordant across overlapping IDPs (Pearson *r* ≈ 0.995; Spearman *ρ* ≈ 0.98), indicating strong robustness to model specification. Detailed results are provided in Supplementary Tables S1–S5.

Among the 3,098 robust IDPs, effects showed clear modality-specific patterns. Median associations were most negative for resting-state fMRI phenotypes (median *β* = −0.0240, *n* = 122) and diffusion-derived phenotypes (dMRI median *β* = −0.0175, *n* = 1,488; DTI-derived median *β* = −0.0207, *n* = 305), with particularly strong negative shifts observed for rs-fMRI network metrics and DTI radial diffusivity. In contrast, structural MRI phenotypes showed modest positive-skewed effects (sMRI median *β* = 0.0095, *n* = 299; sMRI-derived median *β* = 0.0130, *n* = 321).

These patterns were consistent across parcellation schemes, hemispheres (LH median *β* = −0.0157, RH −0.0163), and global measures, arguing against hemisphere-specific artifacts. The strongest individual associations reached |*β*| ≈ 0.054 SD per 1 SD increase in extPRS. Full IDP-level results and group summaries are provided in Supplementary Tables S1–S5.

### Stage 2: extPRS → initiation (direct effect; Cox proportional hazards models)

In Cox proportional hazards models adjusted for baseline covariates (age, sex, ancestry principal components, and site/scanner indicators), higher extPRS was associated with increased risk of substance initiation across all outcomes (*n* = 10,599).

Under the site-clustered standard error specification without SES covariates, each 1 SD increase in extPRS was associated with increased hazards of initiation for alcohol (HR = 1.129, 95% CI 1.076–1.185, *p* = 7.1 × 10^−7^), nicotine (HR = 1.628, 95% CI 1.473– 1.799, *p* = 1.16×10^−21^), cannabis (HR = 1.665, 95% CI 1.447–1.917, *p* = 1.11 × 10^−12^), and any substance (HR = 1.146, 95% CI 1.094– 1.201, *p* = 1.18 × 10^−8^).

Results were robust to alternative variance estimators (HC3) and inclusion of SES covariates, with all hazard ratios remaining greater than 1 and statistically significant. For example, with SES adjustment, hazard ratios were 1.152 (alcohol), 1.562 (nicotine), 1.580 (cannabis), and 1.163 (any substance). These findings indicate that higher externalizing polygenic liability is associated with earlier initiation risk across substances, with the strongest effects observed for nicotine and cannabis. Full results are provided in Supplementary Tables S6–S9.

### Stage 3: extPRS + IDP → initiation (joint Cox models; candidate IDP prioritization)

In Stage 3, we evaluated discovery-significant IDPs from Stage 1 in outcome-specific joint Cox models including both extPRS and a single standardized IDP, and retained associations passing BH-FDR (*q <* 0.05) on the IDP term.

The number of significant IDPs varied substantially across robustness specifications (site-clustered vs. HC3 standard errors; SES covariates on vs. off). Under site-clustered SE without SES covariates, we identified 173 (alcohol), 137 (any substance), 309 (cannabis), and 930 (nicotine) significant IDPs. With SES adjustment, these counts decreased to 31, 32, 137, and 459, respectively.

Overlap analyses indicated moderate replication for alcohol and any substance (Jaccard indices of 0.45 and 0.39, corresponding to 87 and 69 overlapping IDPs), but weaker replication for nicotine (Jaccard = 0.17). Notably, SES-adjusted models largely identified subsets of the non-SES-adjusted results (e.g., 30/31 for alcohol, 31/32 for any substance, 136/137 for cannabis, and ∼451/459 for nicotine), suggesting that SES adjustment primarily reduced statistical power rather than introducing novel signals.

Across all specifications, we identified 1,630 unique outcome–IDP associations, of which 819 (∼50%) replicated in at least two models. Several IDPs were consistently identified across all four specifications, suggesting a core set of robust neuroimaging correlates of substance initiation beyond extPRS.

**Table 1.** Stage 3: Joint Cox models (extPRS + IDP)—number of FDR-significant IDPs by model and outcome. Number of brain imaging-derived phenotypes (IDPs) significantly associated with substance initiation in joint Cox models including externalizing polygenic risk scores (extPRS), after false discovery rate (FDR) correction. Results are shown across two modeling strategies (cluster-robust and HC3 variance estimators) with and without adjustment for socioeconomic status (SES).

| Model group | SES adjustment | Alcohol | Any substance | Cannabis | Nicotine |
| --- | --- | --- | --- | --- | --- |
| cluster | Off | 173 | 137 | 309 | 930 |
| cluster | On | 31 | 32 | 137 | 459 |
| hc3 | Off | 106 | 107 | 0 | 163 |
| hc3 | On | 6 | 47 | 0 | 13 |

### Stage 4: Mediation analysis (ACME/ADE; bootstrap inference)

We evaluated whether brain imaging-derived phenotypes (IDPs) mediate the association between polygenic risk scores (PRS) and substance initiation using bootstrap-based mediation models (5,000 simulations), adjusting for site clustering and socioeconomic status (SES).

Across all outcomes, the direct effect of PRS on initiation (ADE) was strong and highly significant (ADE ≈ 0.04, *p <* 0.001). In contrast, indirect effects through brain IDPs (ACME) were small in magnitude (approximately −3 × 10^−4^ to −6 × 10^−4^), although some reached statistical significance. The proportion mediated was consistently low (*<* 2%), indicating that only a small fraction of genetic risk is transmitted through these brain measures.

Overall, these results indicate that genetic liability to substance initiation operates predominantly through direct pathways, with brain IDPs contributing only minor effects.

#### Outcome-specific mediation results

In the primary model (clustered standard errors with SES adjustment), we observed:

- **Alcohol initiation:** Six IDPs showed significant mediation after multiple testing correction (FDR *<* 0.05). ACME values ranged from −0.00071 to 0.00057, while ADE ranged from 0.0389 to 0.0415. The proportion mediated ranged from −1.77% to 1.44%.
- **Any-substance initiation:** Eleven IDPs remained significant after FDR correction. ACME ranged from −0.00070 to 0.00061, and ADE ranged from 0.0452 to 0.0467. The proportion mediated ranged from −1.53% to 1.31%.
- **Cannabis initiation:** No mediation effects survived multiple testing correction. Nominally significant ACME values ranged from −0.00016 to 0.00019, with ADE between 0.0180 and 0.0202.
- **Nicotine initiation:** No mediation effects survived multiple testing correction. Nominal ACME values ranged from −0.00019 to 0.00027, with ADE between 0.0283 and 0.0300.

These results indicate that robust mediation is present only for alcohol and any-substance initiation, and even in these outcomes, effect sizes are small.

#### Comparison of direct and mediated effects

Across all outcomes, ADE was substantially larger than ACME. ACME estimates were on the order of 10^−4^ and accounted for less than 2% of the total effect.

Visualization of ADE versus ACME (Figure 1) shows a clear separation between large direct effects and minimal mediated effects. This pattern indicates that genetic risk primarily influences substance initiation through direct pathways rather than through baseline brain structure or function.

**Figure 1.**
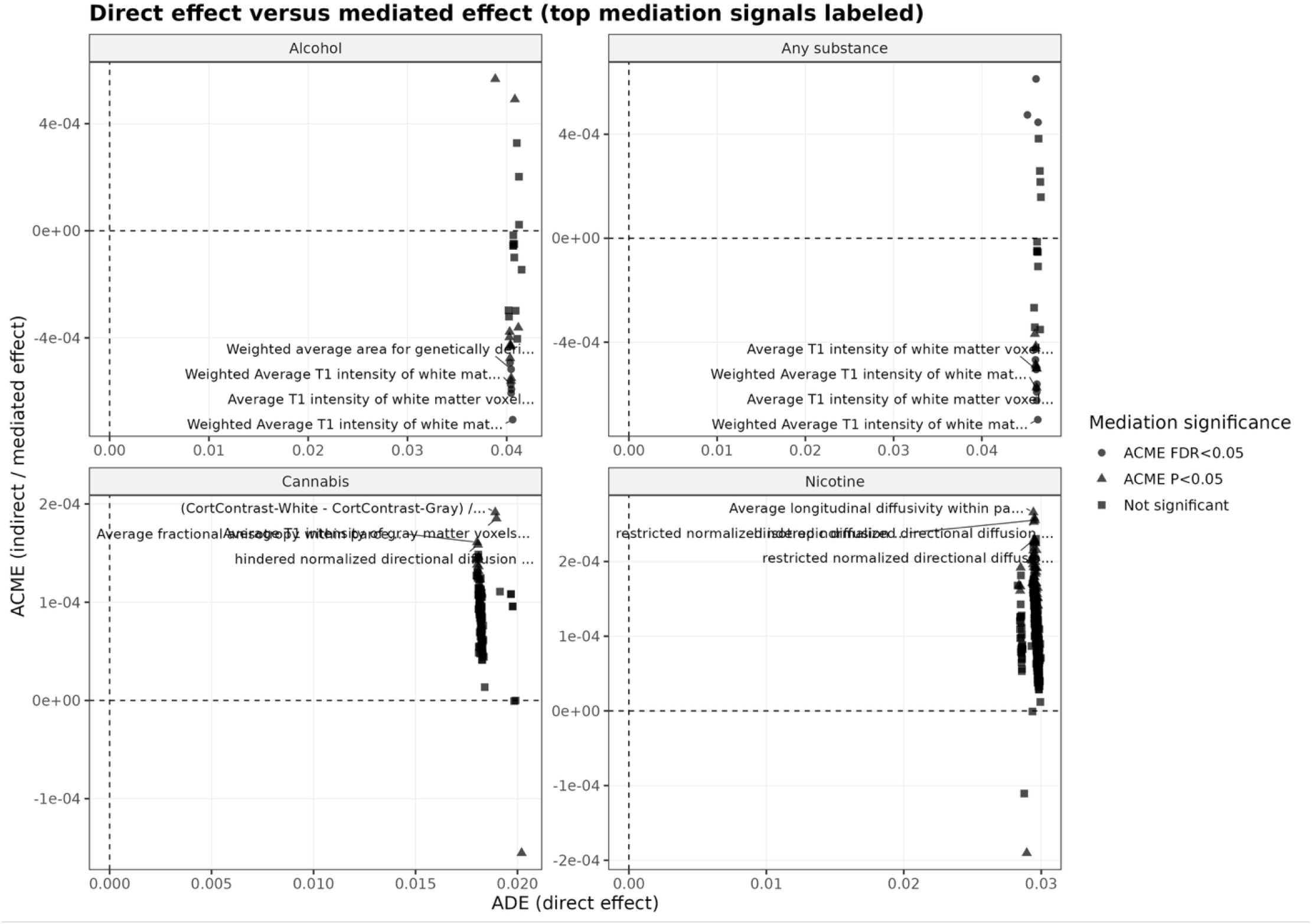
Direct versus mediated effects of externalizing polygenic risk across substance initiation outcomes. Scatter plots show the relationship between the direct effect (ADE; x-axis) and indirect (mediated) effect (ACME; y-axis) for each outcome (alcohol, any substance, cannabis, nicotine). Each point represents a neuroimaging-derived phenotype (IDP). Points are colored by mediation significance (ACME FDR *<* 0.05, ACME *p <* 0.05, or not significant). Across all outcomes, ADE is substantially larger than ACME, with ACME values close to zero (on the order of 10^−4^). This separation indicates that genetic liability primarily influences substance initiation through direct pathways rather than through baseline brain imaging measures. Selected top IDPs are labeled for interpretability.

#### Direction and interpretation of mediation effects

The direction of mediation effects differed by outcome (Figure 2):

**Figure 2.**
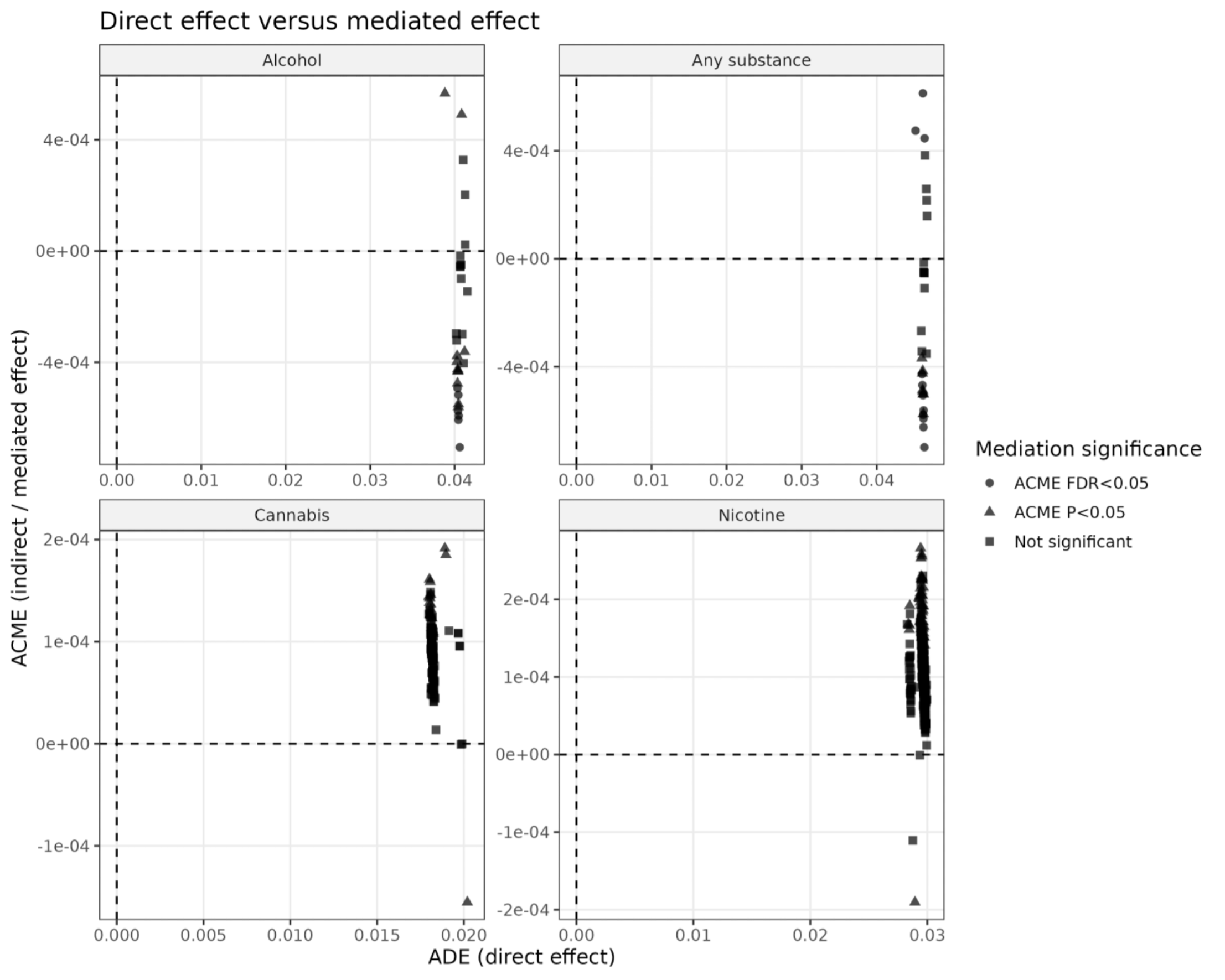
Direction and distribution of mediation effects across outcomes. Scatter plots of ADE (direct effect; x-axis) versus ACME (mediated effect; y-axis) for each outcome without labeling individual IDPs. Alcohol initiation shows predominantly negative ACME values, indicating a consistent buffering (attenuating) effect of brain features on genetic risk. Any-substance initiation shows both positive and negative ACME values, suggesting a mixture of risk-enhancing and protective pathways. Cannabis and nicotine show small and non-significant mediation effects, indicating weak and non-robust mediation patterns.

- **Alcohol initiation:** All significant ACME estimates were negative, indicating that these brain features attenuate the effect of genetic risk, consistent with a buffering or compensatory role.
- **Any-substance initiation:** Both positive and negative mediation effects were observed, indicating the presence of both risk-enhancing and protective pathways.
- **Cannabis and nicotine initiation:** Mediation effects were small and did not survive multiple testing correction, suggesting weak and non-robust mediation.

Across outcomes, the strongest signals involved overlapping brain regions and similar classes of features, including structural measures and diffusion-based metrics, suggesting shared neurodevelopmental pathways.

#### Proportion mediated

Figure 3 summarizes the proportion of genetic risk mediated through brain IDPs. Across all outcomes, the absolute proportion mediated was consistently below 2%:

**Figure 3.**
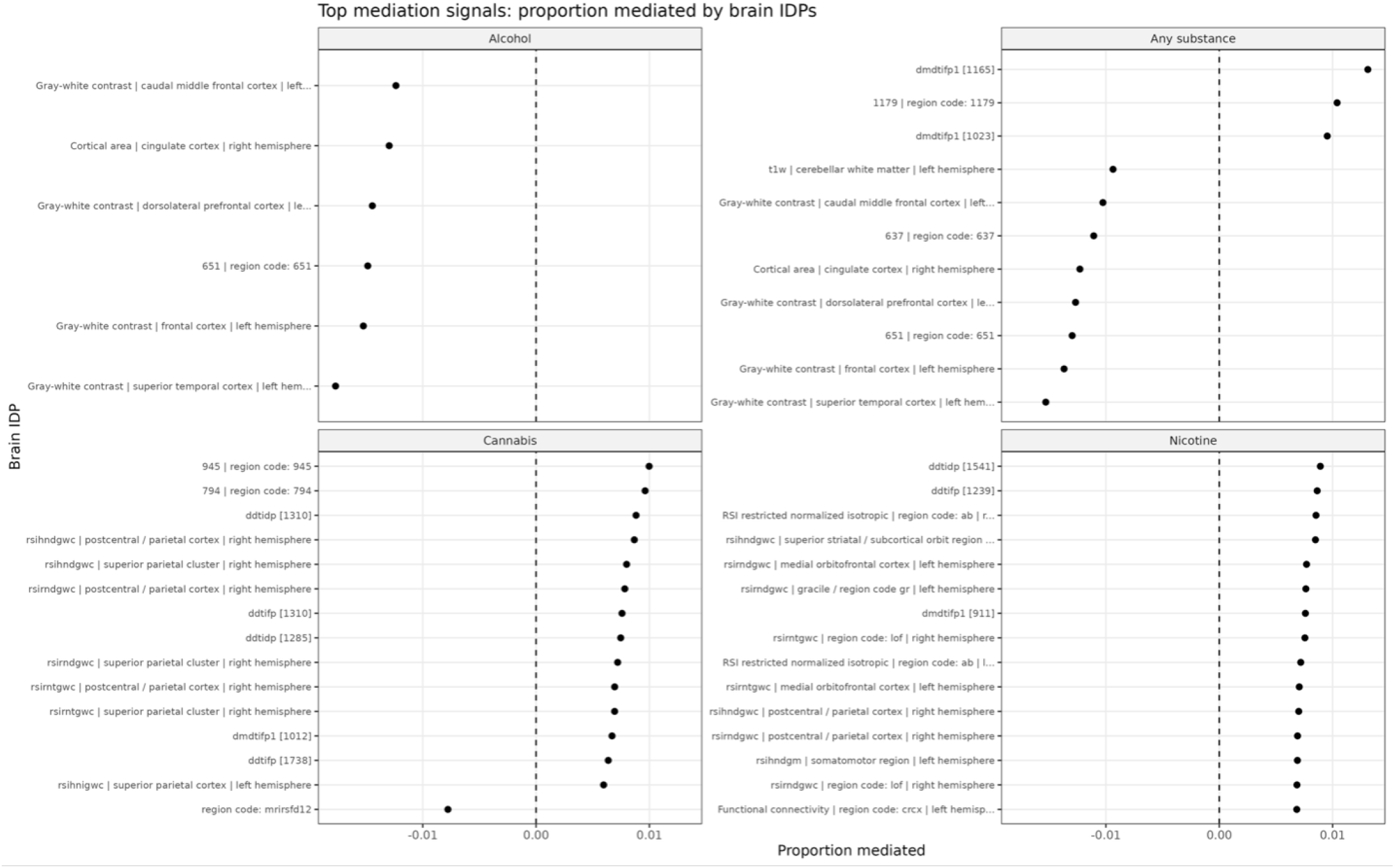
Proportion of genetic risk mediated by brain imaging-derived phenotypes. Top IDPs ranked by proportion mediated for each outcome. Across all outcomes, the absolute proportion mediated is small (generally *<* 2%). Alcohol initiation shows consistently negative mediation effects, indicating a buffering pattern. Any-substance initiation shows mixed positive and negative effects, while cannabis and nicotine show small and non-significant mediation. These results indicate that baseline brain features explain only a minimal proportion of genetic liability.

- **Alcohol:** uniformly negative mediation effects (buffering pattern)
- **Any substance:** mixed positive and negative effects
- **Cannabis and nicotine:** small, non-significant effects

These findings indicate that baseline brain features explain only a minimal proportion of genetic liability.

#### Sensitivity analysis

To assess robustness, we repeated the mediation analysis using heteroskedasticity-consistent (HC3) standard errors while retaining SES adjustment.

The results were consistent with the primary model:

- ADE remained substantially larger than ACME
- Proportion mediated remained below 2%
- Alcohol showed consistently negative mediation effects
- Any substance showed mixed positive and negative effects
- Cannabis and nicotine remained non-significant

Although the specific IDPs identified varied slightly, similar classes of brain features, particularly white matter intensity and diffusion-based measures, were consistently implicated. These results indicate that the observed mediation patterns are robust to alternative variance estimation approaches.

## Discussion

In this study, we developed and applied a multistage framework to link externalizing polygenic liability (extPRS) with baseline neuroimaging phenotypes and prospective substance initiation in the ABCD cohort. Across all stages, we observed a consistent pattern: extPRS is broadly associated with both brain phenotypes and initiation risk, but mediation through baseline brain measures is limited.

### Principal findings

First, extPRS showed widespread associations with neuroimaging phenotypes. Thousands of IDPs were associated with extPRS, and these signals were highly consistent across model specifications. Effect sizes were strongly correlated across sensitivity settings, indicating that the extPRS–IDP associations are stable and not driven by specific modeling choices. These associations showed structured patterns across modalities, with negative shifts in diffusion and functional measures and more modest positive shifts in structural measures.

Second, extPRS was consistently associated with earlier substance initiation across all outcomes. Effect sizes were largest for nicotine and cannabis and more modest for alcohol and any substance, but all associations were statistically significant and robust across variance estimators and SES adjustment. These findings confirm that externalizing genetic liability is a broad risk factor for early substance initiation.

Third, joint models identified a large number of IDPs associated with initiation beyond extPRS. The number of identified IDPs varied across robustness specifications, but overlap analyses revealed a replicated core set of features. Adjustment for SES reduced the number of detectable associations but did not introduce new signals, suggesting that SES primarily affects statistical power rather than altering the underlying relationships. Across models, structural and diffusion-related features were consistently implicated.

Fourth, mediation analyses showed that only a small proportion of the extPRS effect on initiation is mediated through baseline brain phenotypes. Although several IDPs showed statistically significant indirect effects for alcohol and any-substance initiation, these effects were very small in magnitude (ACME on the order of 10^−4^), corresponding to less than 2% of the total effect. For cannabis and nicotine, no mediation effects survived multiple testing correction. Across all outcomes, the direct genetic effect (ADE) was substantially larger than the mediated effect, indicating that genetic liability operates primarily through direct pathways.

### Interpretation

Taken together, these findings suggest that externalizing genetic liability influences substance initiation through multiple pathways, but baseline brain structure and function capture only a small portion of this process.

The widespread extPRS–IDP associations indicate that genetic liability is reflected broadly across brain systems. However, the limited mediation effects suggest that these baseline brain measures are not the primary mechanism linking genetic risk to initiation behavior. Instead, they may represent downstream correlates or parallel processes rather than causal intermediates.

The consistent negative mediation effects observed for alcohol initiation suggest a buffering or compensatory role of certain brain features, particularly structural and white matter measures. In contrast, the mixed positive and negative effects observed for any-substance initiation indicate that both risk-enhancing and protective pathways may be present. The absence of robust mediation for cannabis and nicotine, despite strong direct genetic effects, further supports the conclusion that non-brain pathways or dynamic processes may play a larger role for these outcomes.

One possible explanation is that baseline imaging measures capture only a static snapshot of neurodevelopment, whereas substance initiation is influenced by dynamic processes over time, including environmental exposures, behavioral trajectories, and gene–environment interactions. In this context, the observed IDPs may reflect general neurodevelopmental variation associated with genetic liability rather than specific mediators of initiation risk.

### Methodological implications

This study demonstrates the value of a multistage framework that combines high-dimensional screening, survival modeling, and mediation analysis. By separating discovery (Stage 1), outcome association (Stage 3), and mediation (Stage 4), we reduce the risk of false positives and improve interpretability.

The robustness analyses further strengthen the findings. Results were consistent across different variance estimators and SES adjustment, and a substantial proportion of IDPs replicated across model specifications. This consistency suggests that the identified patterns are not artifacts of modeling choices.

At the same time, the large number of associations in Stage 1 and Stage 3 highlights the importance of careful multiple-testing control and prioritization strategies. The mediation step provides an additional layer of filtering, focusing attention on IDPs with potential mechanistic relevance.

### Limitations

Several limitations should be considered.

First, mediation analyses were based on baseline IDPs, which may not fully capture the dynamic neurodevelopmental processes that influence substance initiation. Longitudinal imaging data may provide stronger tests of mediation.

Second, although we adjusted for a comprehensive set of covariates, residual confounding cannot be ruled out, particularly for environmental and behavioral factors that were not explicitly modeled in the mediation framework.

Third, the mediation models used a simplified structure (extPRS → IDP → initiation), which may not capture more complex pathways involving multiple mediators or interactions.

Fourth, the effect sizes of the mediation signals were small and, although statistically significant in some cases, their practical impact is limited. These findings should therefore be interpreted as indicating limited mediation rather than strong mechanistic pathways.

Finally, the generalizability of the findings may depend on the characteristics of the ABCD cohort and the ancestry composition used in PRS construction.

### Future directions

Future work should extend this framework in several ways:

- Incorporate longitudinal imaging data to evaluate time-varying mediation effects
- Integrate environmental and behavioral factors to test gene– environment pathways
- Apply multivariate or network-based mediation models to capture more complex mechanisms
- Examine developmental trajectories rather than baseline measures alone

## Conclusions

We developed and applied a multistage framework linking externalizing polygenic risk to brain imaging phenotypes and prospective substance initiation in the ABCD cohort.

Across outcomes, extPRS was consistently associated with increased risk of substance initiation, with the strongest effects observed for nicotine and cannabis. ExtPRS was also broadly associated with multimodal neuroimaging phenotypes, indicating widespread neurobiological correlates of genetic liability.

However, mediation analyses showed that only a small proportion (*<* 2%) of the genetic effect is explained by baseline brain measures. Direct genetic effects were substantially larger than mediated effects across all outcomes, indicating that genetic liability operates primarily through direct or non-brain pathways.

Robust mediation signals were observed only for alcohol and any-substance initiation, and even in these cases, effect sizes were small. The identified brain features may play a minor or compensatory role rather than serving as primary mechanisms.

Overall, these findings suggest that while brain phenotypes are associated with genetic liability, they explain only a limited portion of its effect on early substance initiation. This framework provides a scalable and reproducible approach for integrating genetic, neuroimaging, and longitudinal outcome data to prioritize candidate pathways for further investigation.

## Data availability

### Code

The analysis code and scripts used in this study are freely available at the following GitHub repository: https://github.com/mw742/ExtPRS-Brain-SUD.

### Data

This study uses data from the Adolescent Brain Cognitive Development (ABCD) Study (https://abcdstudy.org), held in the NIMH Data Archive (NDA). The ABCD data version used was 5.1. The study is supported by the National Institutes of Health (NIH) and additional federal partners under multiple award numbers, including U01DA041048 and U01DA050987. A full list of funders is available at https://abcdstudy.org/federal-partners.html.

## Author contributions statement

Mengman Wei conceived the study, designed the analytical framework, performed all data processing, statistical analyses, and computational modeling, and drafted the manuscript. All code implementation, data curation, and result interpretation were conducted by Mengman Wei.

Qian Peng provided supervision, general guidance, resource support, and funding acquisition.

## Preprint Notice

This manuscript is a preprint and has not yet undergone peer review. The content is shared to disseminate findings and establish a precedent. Additional analyses and revisions may be incorporated in future versions.

